# Genome-wide mutational signatures of immunological diversification in normal lymphocytes

**DOI:** 10.1101/2021.04.29.441939

**Authors:** Heather E Machado, Emily Mitchell, Nina F Øbro, Kirsten Kübler, Megan Davies, Francesco Maura, Daniel Leongamornlert, Mathijs A. Sanders, Alex Cagan, Craig McDonald, Miriam Belmonte, Mairi S. Shepherd, Robert J. Osborne, Krishnaa Mahbubani, Iñigo Martincorena, Elisa Laurenti, Anthony R Green, Gad Getz, Paz Polak, Kourosh Saeb-Parsy, Daniel J Hodson, David Kent, Peter J Campbell

**Author notes:** Authors contributed equally (Emily Mitchell, Nina F Øbro, Kirsten Kübler). Correspondence to: Peter J Campbell and David Kent.

## Abstract

A lymphocyte suffers many threats to its genome, including programmed mutation during differentiation, antigen-driven proliferation and residency in diverse microenvironments. After developing protocols for single-cell lymphocyte expansions, we sequenced whole genomes from 717 normal naive and memory B and T lymphocytes and hematopoietic stem cells. Lymphocytes carried more point mutations and structural variation than stem cells, accruing at higher rates in T than B cells, attributable to both exogenous and endogenous mutational processes. Ultraviolet light exposure and other sporadic mutational processes generated hundreds to thousands of mutations in some memory lymphocytes. Memory B cells acquired, on average, 18 off-target mutations genome-wide for every one on-target *IGV* mutation during the germinal center reaction. Structural variation was 16-fold higher in lymphocytes than stem cells, with ~15% of deletions being attributable to off-target RAG activity.

**One Sentence Summary:** The mutational landscape of normal lymphocytes chronicles the off-target effects of programmed genome engineering during immunological diversification and the consequences of differentiation, proliferation and residency in diverse microenvironments.

## Main Text

The adaptive immune system depends upon programmed somatic mutation to generate antigen receptor diversity. T lymphocytes use RAG-mediated deletion to generate functional T-cell receptors (TCRs); B lymphocytes also use RAG-mediated deletion to rearrange immunoglobulin (Ig) heavy and light chains, followed by AID-mediated somatic hypermutation and class-switch recombination to further increase diversity (*1–3*). The machinery undertaking this physiological genome editing is tightly regulated, switched on at specific stages of lymphocyte maturation and targeted to specific regions of the genome through chromatin interaction and characteristic sequence motifs.

Off-target activity of proteins generating genomic diversity in lymphocytes can produce mutations in cancer genes with unintended consequences. Whole genome sequencing of malignant lymphoid tumors has revealed high numbers of off-target mutations with signatures resembling those seen at antigen receptors. For example, RAG-mediated deletions at genes regulating B-cell maturation are common in acute lymphoblastic leukemia (*4, 5*); off-target AID-mediated somatic hypermutation is common in diffuse large B-cell lymphoma and chronic lymphocytic leukemia (*6–9*); translocations of oncogenes to immunoglobulin loci occurring during class-switch recombination are common in multiple myeloma (*10*).

It remains unclear whether the burden and signatures of somatic mutations seen in lymphoid cancers would be equivalent in normal lymphocytes, not least because driver mutations in cancers can stall lymphocyte differentiation at stages where physiological genome editing is active (*11, 12*). While single-cell sequencing of 54 B lymphocytes has demonstrated increasing mutation burden with age and evidence of off-target somatic hypermutation (*13*), our novel protocols for producing single-cell derived colonies from naive and memory B and T cells enables the generation of genomes free of whole-genome amplification artifacts, allowing accurate quantification of mutation burdens, signatures and distributions across cells of several lymphocyte subsets.

### Whole genome sequencing of B and T lymphocyte colonies

We and others have shown that growing single hematopoietic stem cells into colonies *in vitro* can enable accurate identification of all classes of somatic mutation using genome sequencing, avoiding the artifacts introduced by whole genome amplification (*14–16*). In order to achieve this for lymphocytes, we developed protocols for expanding flow-sorted single naive and memory B and T lymphocytes *in vitro* to colonies of 30-2000+ cells (**Fig. 1A**; **Methods**).

**Fig. 1.**
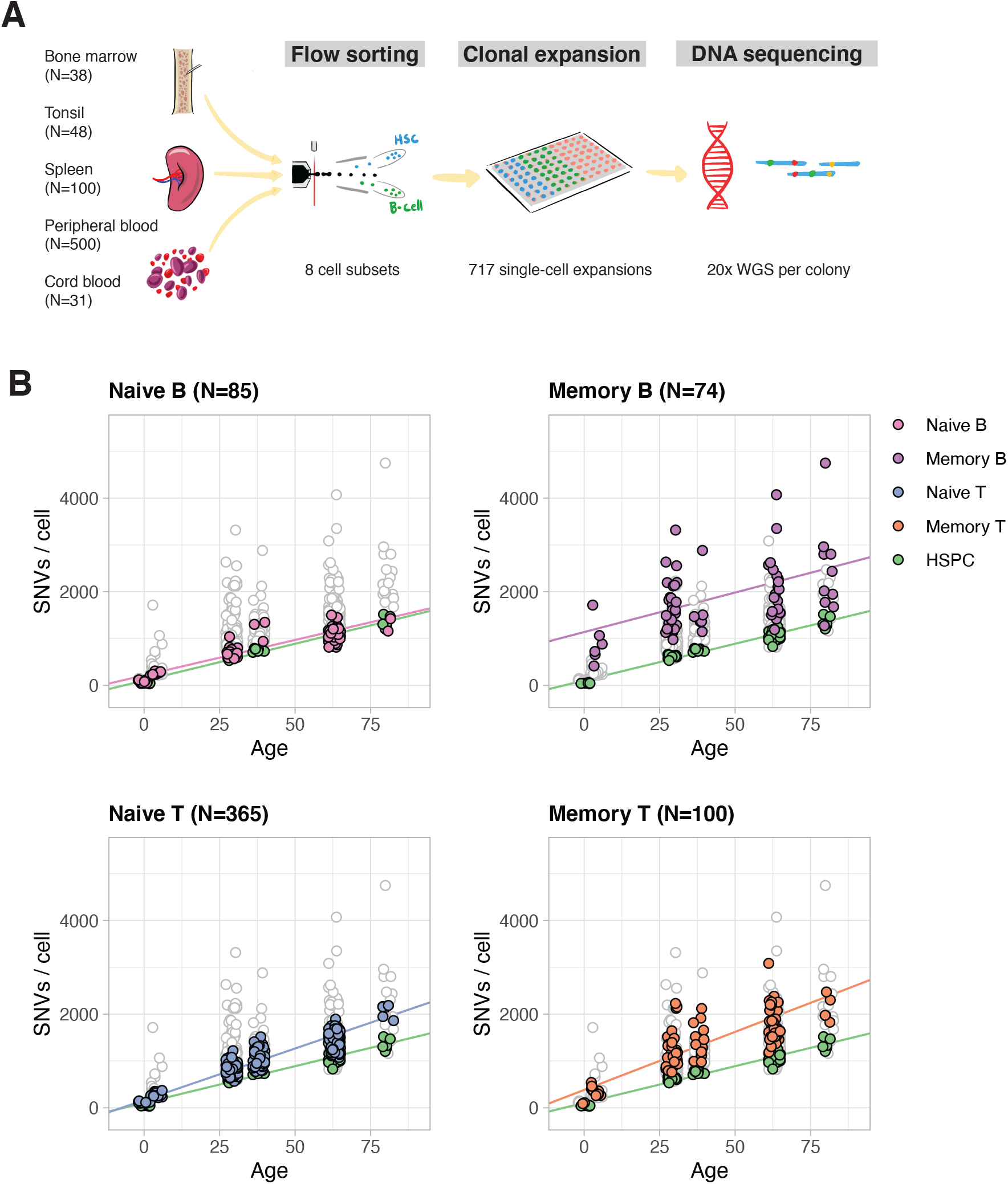
Experimental design and lymphocyte mutation burden with age. (A) Schematic of the experimental design. (B) SNV mutation burden per genome for the four main lymphocyte subsets (colored points), compared with HSPCs (green points). Each panel has all genomes plotted underneath in white with grey outline.

We obtained peripheral blood, spleen and bone marrow samples from four individuals aged 27-81 years, as well as tonsillar tissue from two four-year old children and cord blood from one neonate (**Table S1**). All individuals studied were hematopoietically normal and healthy; one subject had a history of inflammatory bowel disease (Crohn’s disease) treated with azathioprine and the two tonsil donors had a history of tonsillitis. We focused on four main classes of lymphocytes: naive B lymphocytes, memory B lymphocytes, CD4+ and CD8+ naive T lymphocytes, and CD4+ and CD8+ memory T lymphocytes (**Fig. S1**). In one subject we also expanded T-regulatory cells. From five of the subjects, we isolated and expanded hematopoietic stem and progenitor cells (HSPC) to provide a baseline for comparison of mutation burden and signatures in the lymphocytes. We performed whole genome sequencing to an average depth of ~20x, and called somatic mutations using standard, benchmarked bioinformatic algorithms. Average telomere lengths were also estimated from the sequencing data. The final dataset analyzed here comprises 717 whole genomes (**Table S2**).

### High mutation burden of memory lymphocytes and an increased mutation rate in T cells

The overall burden of both single nucleotide variants (SNVs) and insertion/deletions (indels) per cell varied extensively across the dataset, influenced predominantly by age and cell type (**Fig. 1B**), which we quantified with linear mixed effects models. The burden of base substitutions increased linearly with age across all cell types, but the rate of mutation accumulation differed across cell types (p=1×10^−4^ for age-cell type interaction; linear mixed effects model). HSPCs accumulated mutations at ~16 mutations/cell/year (CI_95%_=13-19), similar to previous estimates (*15, 16*). Naive and memory B cells showed broadly similar rates of mutation accumulation (naive B: 15 mutations/cell/year, CI_95%_=12-18; memory B cells: 17 mutations/cell/year, CI_95%_=6-28). T cells, though, had higher mutation rates (naive T: 22 mutations/cell/year, CI_95%_=19-25; memory T cells: 25 mutations/cell/year, CI_95%_=17-32). Overall, this suggests that there are clock-like mutational processes adding mutations at steady rates, with different rates in each lymphocyte subset.

Additionally, there was a significant increase in the burden of base substitutions in lymphocytes that could not be explained by age, especially for memory lymphocytes. Compared to HSPCs, naive B and T lymphocytes had an average of 110 (CI_95%_=5-216) and 59 (CI_95%_=-35-153) extra SNVs per cell, respectively, beyond the effects of age. Memory B and T lymphocytes had an even more pronounced excess of mutations, carrying an average of 1034 (CI_95%_=604-1465) and 277 (CI_95%_=5-549) more mutations than HSPCs respectively. Regulatory T cells were similar to memory T cells in mutation burden. This extra burden of base substitutions presumably represents variants acquired during differentiation: approximately one hundred from HSPC to naive lymphocyte and hundreds to thousands from naive to memory lymphocyte.

While these estimates show that the population average for mutation burden increases with lymphocyte differentiation, we found that the variance in mutation burden across cells also massively increased with differentiation. Thus, compared to a standard deviation of 70 SNVs/cell for HSPCs within a given individual, the values estimated for memory B and T lymphocytes were 820 SNVs/cell and 592 SNVs/cell respectively (p<10^−16^ for heterogeneity of variance across cell types).

Indels showed similar patterns, accumulating at an average of 0.7/cell/year in HSPCs (CI_95%_=0.5-0.9; p=0.02 for age-cell type interaction) with lymphocytes, especially memory B and T cells, carrying an excess burden compared to hematopoietic stem cells (**Fig. S2**).

Driver mutations, defined as those under positive selection, are frequently present in ageing normal tissues (*17–20*), including blood (*21–25*). We did not observe any significant enrichment of non-synonymous variants in driver genes associated with age-related clonal hematopoiesis or lymphoma in our normal lymphocytes, however.

### Mutational signatures in B and T lymphocytes are distinct from HSPCs

In order to determine whether the excess mutations observed in lymphocyte subsets were due to a specific mutational process, we extracted mutational signatures across lymphocyte compartments (**Fig. 2**). Like HSPCs, the vast majority of mutations in naive B and T cells were derived from two mutational signatures. One of these, SBS1, is caused by spontaneous deamination of methylated cytosines (*26, 27*), and accounted for 14% of mutations in HSPCs and naive B and T lymphocytes. Nearly all remaining somatic mutations in these cellular compartments had the typical signature of endogenous mutations in HSPCs (*15, 16*), which we term ‘SBSblood’ (**Fig. S3**). The burden of both signatures correlated linearly with age (**Fig. S4**), suggesting that they represent clock-like endogenous mutational processes.

**Fig. 2.**
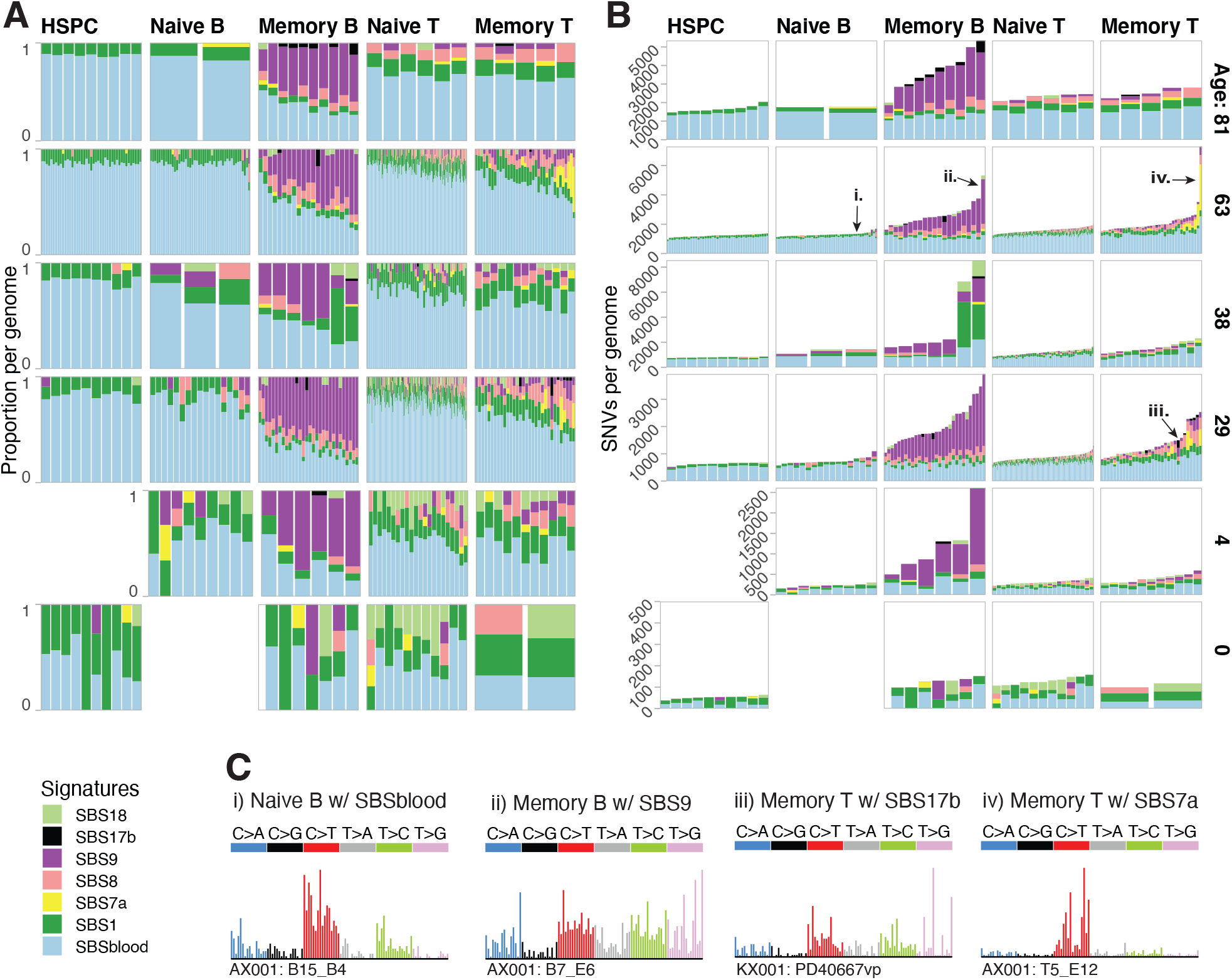
Mutational processes in lymphocytes. (A) The proportion of SNVs and (B) SNV burden per mutational signature. Each column represents one genome. Signatures are identified by the programs *hdp* and *sigprofiler* and attribution per genome is performed by *sigfit*. Per genome, signatures with a 90% CI lower bound of less than 1% are excluded from plotting. (C) Mutational spectra of single genomes enriched in the specified mutational signature. The specific genome plotted is identified with the corresponding roman numeral in panel (B). The trinucleotide contexts of the mutations, which compose the x-axis, are ordered as in Fig. S3.

For memory B and T lymphocytes, the absolute numbers of mutations attributed to these two endogenous signatures were broadly similar to those seen in naive B and T lymphocytes (**Fig. 2**). The hundreds to thousands of extra mutations seen in memory B and T lymphocytes derived from additional mutational signatures: SBS7a, SBS8, SBS9, and SBS17b. While signatures SBS8 and SBS9 show correlations with age, SBS7 and SBS17a do not, consistent with them being sporadic (**Fig. S4**). SBS7a and SBS17b likely represent exogenous mutational processes, discussed in the next section, while SBS9 is differentiation-associated, discussed thereafter.

### Mutational signatures as clues to historical tissue residency of circulating lymphocytes

SBS7 is the canonical signature of ultraviolet light damage, the predominant mutational process in melanoma (*28*), normal skin (*18, 29*) and mycosis fungoides (*30, 31*), a T-cell lymphoma derived from skin-resident memory T cells (*32*). The signature we extracted in memory lymphocytes matches the features of SBS7, with a predominance of C>T substitutions in a dipyrimidine context, transcriptional strand bias and a high rate of CC>TT dinucleotide substitutions (**Fig. S5**). We found a substantial contribution of SBS7 (>10% of mutations; mean=757/cell, range 205-2783) and CC>TT dinucleotide substitutions in 9/100 memory T cells. Interestingly, memory lymphocytes with high SBS7 had significantly shorter telomeres than other memory T cells (p=0.01, Fisher’s method; **Fig. S6**), indicative of increased proliferation. UVB radiation, the most mutagenic, only penetrates human skin to a depth of 10-50μm (*33*), which suggests that these memory T lymphocytes were skin-resident at some stage during their life. That such a high fraction (9/100) of memory T lymphocytes from peripheral blood exhibit a skin-specific mutational process suggests that skin-resident T lymphocytes represent a large and dynamic population, frequently recirculating via the blood system (*34, 35*).

A second unexpected signature in memory lymphocytes was SBS17. This signature has been observed in cancers of the stomach and esophagus (*36*) and occasionally in B (*36*) and T cell lymphomas (*37*). This signature, characterized by T>G mutations in a TpT context, accounted for >10% of mutations (4SD above mean) in 3/74 memory B and 1/100 memory T lymphocytes. SBS17 has been linked to 5-flurouracil chemotherapy in metastatic cancers (*38, 39*), but its occurrence in esophageal and gastric cancers (as well as our samples here) is independent of treatment. If its incidence in upper gastrointestinal tract cancers is caused by some unknown local mutagen, then the presence of SBS17 in memory lymphocytes may again represent evidence of a specific microenvironmental exposure associated with tissue residency in the gastric and/or esophageal mucosa.

### Signatures of the germinal center reaction

The extensive variation in memory B cell mutation burden is primarily explained by mutational signature SBS9, which accounts for 42% of mutations (mean, 780 mutations/cell), at times tripling the baseline mutation burden. This signature is characterized by mutations at A:T base-pairs, especially T>G in a TpW context, and has been found in both healthy (*13*) and malignant post-germinal center B cells (*6–9*). As reported for lymphoid malignancies (*8, 40, 41*), we found that SBS9 has a different spectrum to that of somatic hypermutation at immunoglobulin loci (**Fig. 3A**) and that the two mutational signatures have very different distributions across the genome (**Fig. S7**).

**Fig. 3.**
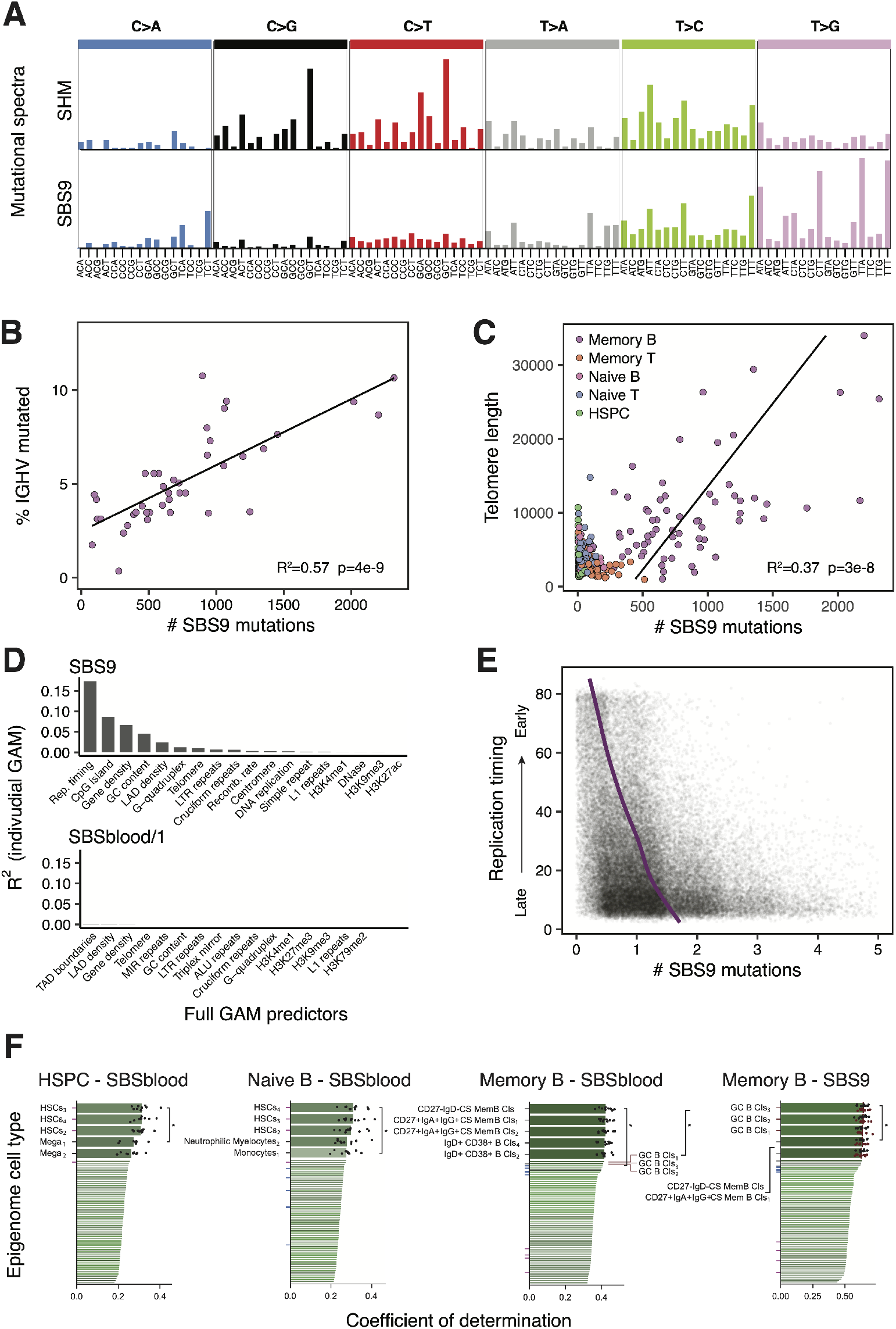
Correlation of SBS9 with genomic attributes and timing of mutational processes. (A) Mutational spectra of the SBS9 signature and the SHM signature, the latter identified independently by de-novo extraction from 1MB bins of memory B cell mutations. (B) Correlation of SBS9 and the extent of SHM as measured by the proportion of the IGHV mutated in the productive rearrangement of memory B cells. (C) Correlation of SBS9 and telomere length per genome. The regression line is for memory B cells. (D) Explanatory power of each genomic feature found significant in the full GAM model (expressed as the R^2^ of the individual GAM model) for predicting number of SBS9 mutations (top) or number of SBSblood/1 mutations per 10Kb window. (E) Replication timing and the number of SBS9 mutations per 10Kb window. The purple line is the GAM regression prediction. The x-axis is truncated at 5, excluding 0.3% of the data. Points have random noise (−0.5 to 0.5) on the SBS9 mutation measurement to facilitate visualization. (F) Performance of prediction of genome-wide mutational profiles (number of mutations indicated) attributable to particular mutational signatures from histone marks of 149 epigenomes representing distinct blood cell types and different phases of development (subscripts indicate replicates); ticks are colored according to the epigenetic cell type (purple, HSC; blue, naive B cell; grey, memory B cell; maroon, GC B cell); black points depict values from ten-fold cross validation; p-values were obtained for the comparison of the 10-fold cross validation values using the two-sided Wilcoxon test (Cls, cells; CS, class switched; GC, germinal center; HSC, hematopoietic stem cell; Mem, memory). To compare signatures in memory B cells, we trained models on 33,950 mutations with the highest SBS9 probability (maroon dots) and found that the models using germinal center B cells epigenomes were significantly better than for an equally sized set of SBSblood mutations (p=1.1×10^−5^). Mega: megakaryocyte.

In our normal memory B lymphocytes, the number of SBS9 mutations genome-wide showed a strong linear correlation with the number of mutations in the productive V(D)J rearrangement of the *IGHV* gene, despite their different spectra (**Fig. 3B**). Strikingly, 57% of the variation in the number of SBS9 mutations in memory B cells genome-wide could be explained by the number of mutations in *IGHV* (R^2^=0.57, p=4×10^−9^, linear regression). The density of mutations was 270,000-fold greater at the *IGHV* locus than for SBS9 mutations genome-wide, confirming the precise targeting of somatic hypermutation to antibody regions. Nonetheless, the genome is large, and even this high degree of mutational targeting means that every 1 on-target IGV mutation is accompanied by an average of 18 SBS9 mutations elsewhere in the genome.

Another feature of the germinal center reaction is increased telomerase activity in B cells (*42, 43*). We therefore estimated telomere lengths from the genome sequencing data for our dataset. Whereas HSPCs and other lymphocyte subsets showed tightly clustered telomere lengths and the expected decrease with aging, telomere lengths in memory B cells were longer, more variable and actually increased with age (excluding tonsil samples; R^2^=0.13, p=3×10^−3^, linear regression; **Fig. S8**). Telomere lengths also correlated linearly with the number of SBS9 mutations genomewide (excluding tonsil samples; R^2^=0.37, p=3×10^−8^, linear regression; **Fig. 3C**). These data confirm that telomeres do lengthen in the germinal center reaction, and provide further evidence that off-target SBS9 mutations are generated in the germinal center.

### A replicative-stress model of SBS9 mutation

The cytosine deaminase AID initiates on-target somatic hypermutation at immunoglobulin loci, which generates damage (and consequent mutation) at C:G base-pairs. On-target mutations at A:T base-pairs during SHM arise through errors introduced during translesion bypass of AID-deaminated cytosines by polymerase-eta (*44–48*), which has an error spectrum weighted towards a TpW context (*49, 50*). As has been noted in lymphoid malignancies (*8, 36, 44, 51*), off-target SBS9 has a different spectrum from on-target, AID-mediated somatic hypermutation, something we also observe in normal lymphocytes. In particular, SBS9 has a paucity of mutations at C:G base-pairs and an enrichment of T mutations in TpW context (**Fig. 3A**), which makes the role of AID unclear. The genome-wide distribution of off-target AID-induced deamination has been measured directly (*52*), and shows a predilection for highly transcribed regions with active chromatin marks, which tend to be early-replicating.

To explore whether genomic regions with high SBS9 burden show the same distribution, we used general additive models to predict SBS9 burden from 36 genomic features, including gene density, chromatin marks and replication timing across 10kb genome bins. After model selection, 18 features were included in the regression (R^2^=0.20; **Fig. 3D**, **Table S3**). Replication timing is by far the strongest predictor, with increased mutation density in late-replicating regions, individually accounting for 17% of the variation in the genomic distribution of SBS9 (**Fig. 3E**). In contrast, replication timing accounted for only 0.6% of variation in density of SBSblood/SBS1 mutations in memory B cells and 0.1% in HSPCs. The next 4 strongest predictors of SBS9 distribution were all broadly related to inactive versus active regions of the genome (distance from CpG islands, gene density, GC content, and LAD density: individual R^2^ 0.09, 0.07, 0.05, and 0.02, respectively). For each variable, mutation density increased in the direction of less active genomic regions, in contrast to AID-induced deamination, which occurs in actively transcribed regions (*52*).

Taken together, our data demonstrate that SBS9 accumulates during the germinal center reaction, evidenced by its tight correlation with both on-target SHM and telomere lengthening. However, the relative sparsity of mutations at C:G base-pairs and the distribution of SBS9 to late-replicating, repressed regions of the genome make it difficult to argue that AID is involved. Instead, we hypothesize that SBS9 arises from polymerase-eta bypass of other background DNA lesions induced by the high levels of replicative and oxidative stress experienced by germinal center B cells. Normally, mismatch repair and other pathways would accurately correct such lesions, but the high expression of polymerase-eta in germinal center cells (*53*) provides the opportunity for error-prone translesion bypass to compete. The enrichment of SBS9 in late-replicating, gene-poor, repressed regions of the genome, regions where mismatch repair is typically less active (*54, 55*), would be consistent with this as a model of SBS9 mutation. The shift from T>C transitions seen in the SHM signature (*36, 44, 56*) and polymerase eta in vitro spectrum (*49, 50*), to T>G transversions in SBS9 (**Fig. 3A**) may be related to the strong transversion bias observed in late replication (*57*).

### Association with cell-type-specific epigenetic marks reveals timing of mutational processes

Among human cell types, lymphocytes are unusual for passing through functionally distinct, long-lived differentiation stages with on-going proliferative potential. Since variation in mutation density across the genome is shaped by chromatin state, a cell’s specific distribution of somatic mutation provides a record of the past epigenetic landscape of its ancestors back to the fertilized egg (*58–60*). We thus hypothesized that the distribution of clock-like signatures will inform on the cell types present in a given cell’s ancestral line-of-descent. In contrast, the distribution of sporadic or episodic signatures can inform on the differentiation stage exposed to that particular mutational process.

We used Random Forest regression to model the distribution of somatic mutations across the genome compared to 149 epigenomes representing 48 distinct blood cell types and differentiation stages (*61–63*). We found that mutations resulting from the clock-like signature SBSblood in HSPCs correlated best with histone marks from hematopoietic stem cells (p=0.002, Wilcoxon test; **Fig. 3F**), consistent with mutation accumulation in undifferentiated cells. Notably, SBSblood mutational profiles in naive B cells also correlated better with the epigenomes of hematopoietic stem cells than naive B cells (p=0.004; **Fig. 3F**). This implies that the majority of SBSblood mutations in naive B cells were acquired pre-differentiation, consistent with on-going production of these cells from the HSPC compartment throughout life and a relatively short-lived naive B differentiation state. In contrast, SBSblood mutations in naive T cells mapped best to the epigenomes of CCR7^+^/CD45RO^−^/CD25^−^/CD235^−^ naive T cells (p=0.049; **Fig. S9**), consistent with a long-lived, numerically predominant pool of naive T cells generated in the thymus during early life. For memory B cells, SBSblood most closely correlated with histone marks from that cell type and not earlier differentiation stages (p=0.02; **Fig. 3F**), suggesting that the majority of their lineage has been spent as a memory B cell.

For the sporadic mutational processes, SBS9 mutations most closely correlated with germinal center B cell epigenomes (p=0.049; **Fig. 3F**). This is consistent with our finding of a correlation between SBS9 and other germinal center-associated processes (SHM and telomere lengthening), providing further evidence that SBS9 arises as a by-product of the germinal center reaction. For SBS7, the signature of ultraviolet light exposure seen in memory T cells, the genomic distribution more tightly associated with epigenomes of differentiated T cells than naive T cells (**Fig. S9**), supporting the hypothesis that SBS7 mutations accumulate in differentiated T cells.

### V(D)J recombination (but not CSR) machinery is associated with off-target structural variants

Both V(D)J recombination and class-switch recombination (CSR) generate large deletions in the Ig/TCR gene regions during lymphocyte maturation. This programmed genome engineering is associated with off-target structural variation (SV) in human lymphoid malignancies (*4, 5, 64*) and murine cell models (*65*), and genomic analysis of associated binding motifs suggests substantial mutagenic potential (*66*); however, rates and patterns of SVs have not been studied in normal human lymphocytes.

We found 1037 SVs across 635 lymphocytes, of which 85% occurred in Ig/TCR regions, consistent with 1-2 V(D)J recombination events per lymphocyte and 0-2 class-switch recombination events per memory B cell. Excluding Ig/TCR gene regions, B and T lymphocytes carried more SVs than HSPCs, with 103/609 (17%) of lymphocytes having at least one off-target SV (compared to a single SV in 82 HSPCs; p=9×10^−5^, Fisher exact test). Memory B and T lymphocytes had higher non-Ig/TCR SV burdens than their respective naive subsets (27% memory B versus 5% naive B cells; 25% memory T versus 15% naive T cells; p=1×10^−5^). Although we saw occasional instances of more complex abnormalities, including chromoplexy (*67*) (**Fig. 4A**) and cycles of templated insertions (*64*), most non-Ig/TCR SVs were deletions (49%), several of which affected genes mutated in lymphoid malignancies (**Fig. 4B**).

**Fig. 4.**
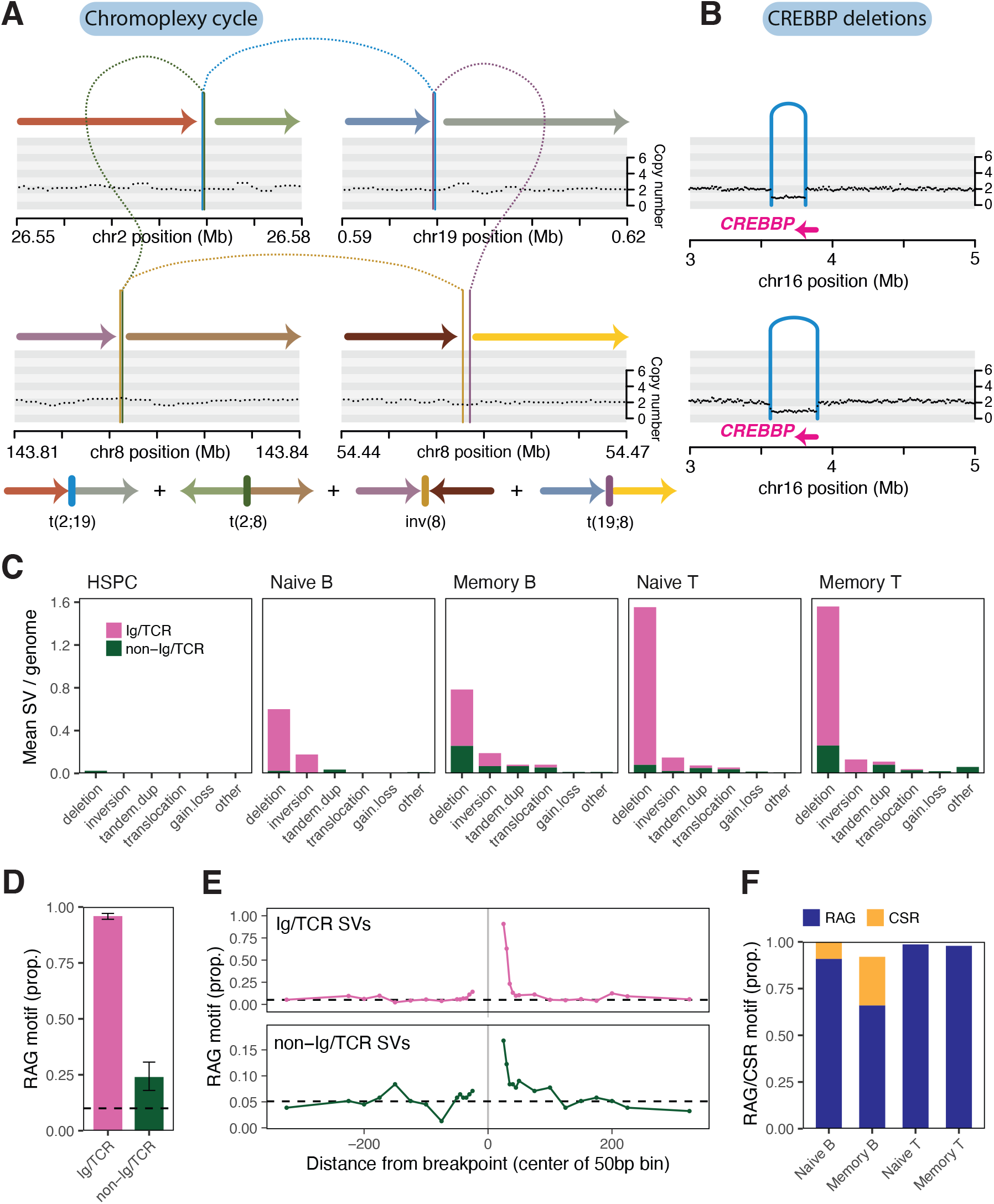
Structural variation burden and off-target RAG-mediated deletion. (A) Chromoplexy cycle (sample PD40667sl, donor KX002). (B) CREBBP deletions (samples PD40521po, donor KX001 and BMH1_PlateB1_E2, donor AX001). (C) Burden of structural variants per cell type. (D) The proportion of deletions with an RSS (RAG) motif within 50bp of the breakpoint. The black dashed line represents the genomic background rate of RAG motifs. Error bars represent 95% bootstrap confidence intervals. (E) The proportion of deletions with an RSS (RAG) motif as a function of distance from the breakpoint, with a positive distance representing bases interior to the deletion, and a negative value representing bases exterior to the breakpoint. The black dashed line represents the genomic background rate of RAG motifs. (F) Proportion of deletions with an RSS (RAG) or switch (CSR) motif.

V(D)J recombination is mediated by RAG1 and RAG2 cutting at an ‘RSS’ DNA motif comprising a heptamer and nonamer with intervening spacer. 24% of non-Ig/TCR and 96% of Ig/TCR SVs had a full RSS motif or the heptamer within 50bp of a breakpoint (**Fig. 4C-D**). Taking into account the baseline occurrence of these motifs using genomic controls, we estimate that 12% of non-Ig/TCR and 84% of Ig/TCR SVs were RAG-mediated, especially deletions (~15% of non-Ig/TCR deletions). As expected, the RSS motif was typically internal to the breakpoint (62% and 91% for non-Ig/TCR and Ig/TCR SVs). We observed a rapid decay in the enrichment of RAG motifs with distance from breakpoints after the first 50bp window, reaching background levels by ~100bp (**Fig. 4E**). During V(D)J recombination, the TdT protein adds random nucleotides at the dsDNA breaks - this also seems to occur in off-target SVs, with RAG-mediated events enriched for insertions of non-templated sequence at the breakpoint (44% and 88% for non-Ig/TCR and Ig/TCR SVs, respectively, versus 21% of off-target SVs without RSS motif; p=9×10^−3^, Fisher exact test).

CSR is achieved through AID cytosine deamination at WGCW clusters (*68, 69*), deleting IgH constant region genes and changing the antibody isotype. CSR accounted for the majority of Ig SVs not attributed to RAG; together, RAG and AID accounted for 92% Ig SVs in memory B cells and 100% in naive B cells (**Fig. 4F**). In contrast, none of the non-Ig/TCR SVs had CSR AID motif clusters, suggesting that class-switch recombination is exquisitely targeted.

### Comparison with malignancy

A long-standing controversy in cancer modelling is whether tumors require additional mutational processes to acquire sufficient driver mutations for oncogenic transformation (*70*). In many solid tissues, cancers do have higher mutation burdens than normal cells from the same organ (*20, 71, 72*), but myeloid leukemias do not (*14*). To address this question in lymphoid malignancies, we compared our normal B and T lymphocytes to 7 blood cancers (*51, 73*). SNV burdens for follicular lymphoma, DLBC lymphoma and multiple myeloma were considerably higher than normal lymphocytes (**Fig. 5A-B**). In contrast, point mutation burdens observed in Burkitt lymphoma, mutated CLL, unmutated CLL and AML were well within the range of normal lymphocytes. In contrast, all lymphoid malignancies showed higher rates of SV than normal cells.

**Fig. 5.**
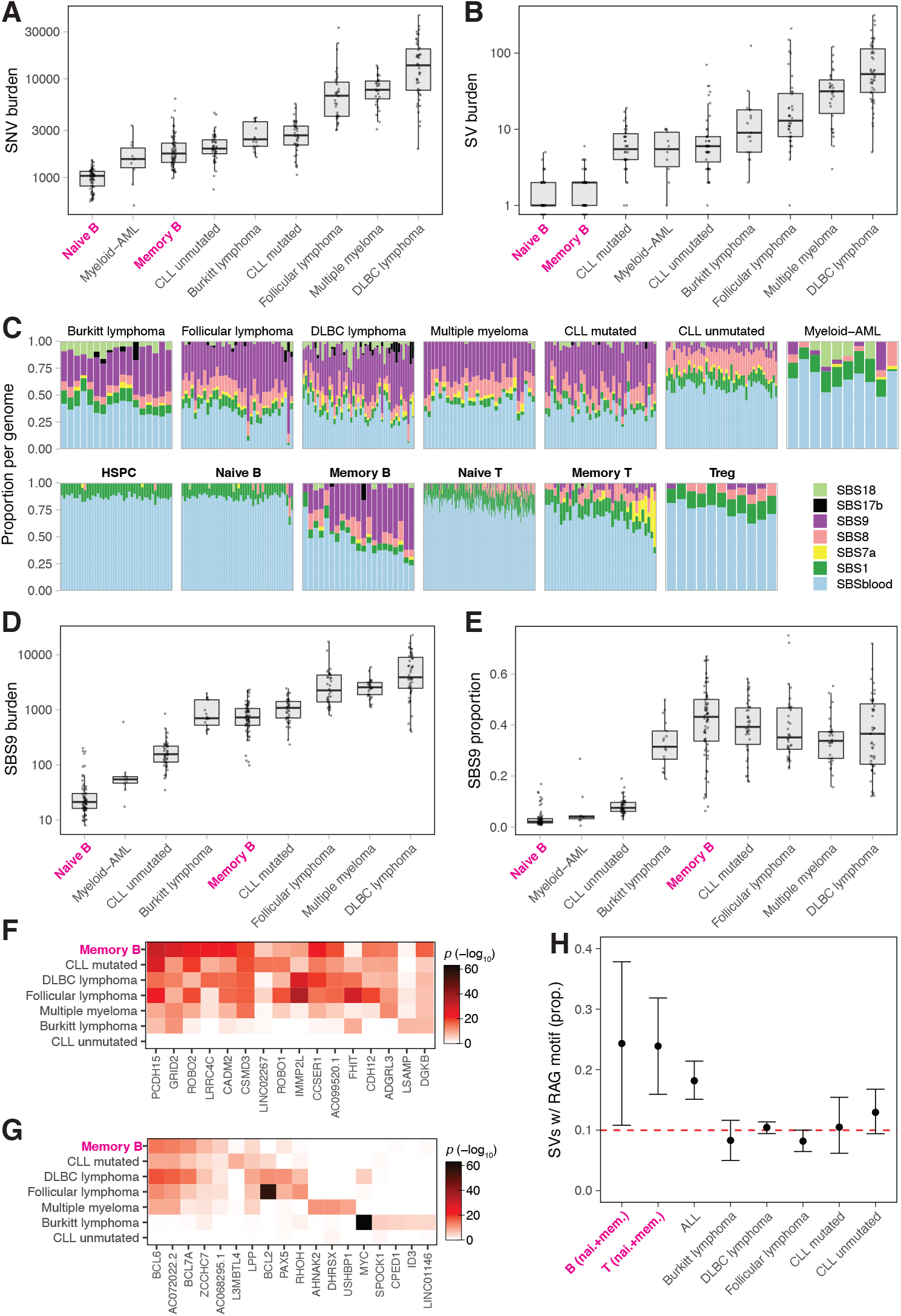
Comparison of mutational patterns with malignancy. (A) SNV and (B) SV burden by cell/malignancy type. Boxes show the interquartile range and the center horizontal lines show the median. Each genome is plotted as one point. (C) Proportion of mutational signatures per genome. Signatures are identified by the programs *hdp* and *sigprofilier* and attribution per genome is performed by *sigfit.* Per genome, signatures with a 90% CI lower bound of less than 1% are excluded from plotting. (D) SBS9 burden and (E) proportion by cell/malignancy type. (F) SBS9 and G) SHM signature enrichment per gene. (H) Proportion of SV’s with RSS (RAG) motifs within 50bp of a breakpoint. (A-E,H) Normal lymphocytes (bold) exclude pediatric samples.

The elevated point mutation burden could arise from increased activity of mutational processes already present in normal cells, or the emergence of distinct, cancer-specific mutational processes. The vast majority of mutations present across all B-cell malignancies could be attributed to the same mutational processes active in normal memory B cells, and at broadly similar proportions (**Fig. 5C-E**). This suggests that the elevated SNV mutation burdens of DLBC lymphoma, follicular lymphoma and multiple myeloma are due to globally increased activity of the same mutational processes active in normal lymphocytes, rather than enrichment of any single process or novel, cancer-restricted DNA repair defects. This is different to, say, colorectal cancer or breast cancer, where the elevated mutation burden is caused by emergence of cancer-specific mutational processes (*72, 74*).

A feature of somatic mutations in B-cell lymphomas is clustering of off-target somatic hypermutation in highly expressed genes. For both SBS9 (**Fig. 5F**) and off-target SHM mutations (**Fig. 5G**), we found considerable overlap in genes with elevated mutation rates. For example, *BCL6, BCL7A* and *PAX5* had enrichment of mutations with the SHM signature in both normal and post-germinal malignant lymphocytes. Likewise, of the 100 genes most enriched for SBS9 in normal memory B cells, 64% were also SBS9-enriched (top 1%) in ≥3 of the 5 post-germinal malignancies.

About 10% of normal lymphocytes have a non-Ig/TCR RAG-mediated SV, accounting for 24% of off-target rearrangements. Across lymphoid malignancies, acute lymphoblastic leukemias (*4*) had similarly high proportions of RAG-mediated events, but in much higher numbers, as reported previously (*4, 5*) (**Fig. 5H**). For other lymphoid malignancies, although the proportions were low, the absolute numbers of RAG-mediated SVs (≥0.5/lymphoma) were broadly comparable to those seen in normal lymphocytes (**Fig. S10**). This suggests that malignant transformation of lymphocytes is associated with the emergence of cancer-specific genomic instability, generating a genome with considerably more large-scale rearrangement.

### Conclusions

Unique among human cell types, a lymphocyte experiences long periods of its life in diverse microenvironments, be it bone marrow, thymus, lymph node, skin or mucosa. These stages are interspersed with short-lived bursts of differentiation, each of which is associated with proliferation and/or programmed genome engineering to improve antigen recognition. Our data show that each phase of lymphocyte differentiation contributes an additional burden of mutations - naive B and T lymphocytes have ~100 more mutations than hematopoietic stem cells; memory lymphocytes have hundreds to thousands more than naive lymphocytes. The signatures of these mutations reflect both the unintended by-products of immunological diversification and exposure to exogenous mutagens; their genomic distribution reflects the chromatin landscape of the cell at the time the mutational process was active. The rare lymphocyte that transforms to cancer draws its stock of somatic mutations from the same mutational processes that are active in normal lymphocytes.

## Code availability

An exhaustive repository of code for statistical analyses reported in this manuscript is available at https://github.com/machadoheather/lymphocyte_somatic_mutation

## Acknowledgments

The authors would like to thank Federico Abascal, Tim Coorens, Timothy Butler and Simon Brunner for valuable guidance in data analysis, the CASM lab, including Laura O’Neill and Calli Latimer, for sample and data management, and CASM IT for technical support. This research was supported by the Cambridge NIHR BRC Cell Phenotyping Hub and staff, including Esther Perez and Natalia Savinykh who provided advice and support in flow cytometry and cell sorting. We are especially grateful to the tissue donors and their families and to the Cambridge Biorepository for Translational Medicine for the gift of tissue from transplant organ donors.

## Materials and Methods

### Samples

Human blood mononuclear cells (MNCs) were obtained from four sources: 1) bone marrow, spleen and peripheral blood from three transplant organ donors (KX001, KX002, KX003) recruited from Cambridge University Hospitals NHS Trust, Addenbrooke’s Hospital (by Cambridge Biorepository for Translational Medicine, Research Ethics Committee approval 15/EE/0152), 2) peripheral blood from one patient (AX001) recruited from Addenbrooke’s Hospital (approval 07-MRE05-44), 3) tonsil from two patients (TX001, TX002) recruited from Addenbrooke’s Hospital (approval 07-MRE05-44), and 4) one cord blood (CB001) collected with informed consent by StemCell Technologies (catalog #70007) (Table S1). All sources were hematopoietically normal and healthy. Donor KX002 had a history of Crohn’s disease and treatment with Azathioprine. Patients TX001 and TX002 had a history of tonsillitis. MNCs from (1), (2) and (3) were extracted using Lymphoprep (Axis-Shield), depleted of red blood cells using RBC lysis buffer (BioLegend) and frozen viable in 10% DMSO. Cord blood MNCs (4) were received frozen and then CD34^+^ selected using the EasySep human whole blood CD34 positive selection kit (Stemcell Technologies) as per the manufacturer’s instructions, with the CD34^+^ fraction used for hematopoietic stem and progenitor cell (HSPC) cultures and the CD34^−^ fraction used for lymphocyte cultures. Additional peripheral blood MNCs from (1) also underwent CD34 positive selection and was used for HSPC cultures.

### Flow cytometry

MNC samples were sorted by flow cytometry at the NIHR Cambridge BRC Cell Phenotyping Hub on AriaIII or AriaFusion cell sorters into naive B lymphocytes (CD3^−^CD19^+^CD20^+^CD27^−^CD38^−^IgD^+^), memory B lymphocytes (CD3^−^CD19^+^CD20^+^CD27^+^CD38^−^IgD^−^), naive T lymphocytes (CD3^+^CD4/CD8^+^CCR7^+^CD45RA^high^), memory T lymphocytes (CD3^+^CD4/CD8^+^CD45RA^−^), regulatory T cells (Tregs: CD3^+^CD4^+^CD25^high^CD127^−^) and HSPCs (CD3^−^CD19^−^CD34^+^CD38^−^CD90^+^CD45RA^−^) (Fig. S1). HSPCs from AX001 included HSCs (CD34^+^CD38^−^) and progenitors (CD34^+^CD38^+^CD10^−/dim^). The antibody panels used are as follows: lymphocytes (excluding Tregs): CD3-APC, CD4-BV785, CD8-BV650, CD14-BV605, CD19-AF700, CD20-PEDazzle, CD27-BV421, CD34-APC-Cy7, CD38-FITC, CD45RA-PerCP-Cy5.5, CD56-PE, CCR7-BV711, IgD-PECy7, Zombie-Aqua; Tregs: CD3-APC, CD4-BV785, CD8-BV650, CD19-APC-Cy7, CD45RA-PerCP-Cy5.5, CD56-PE, CCR7-FITC, CD25-PECy5, CD127-PECy7, CD69-AF700, CD103-BV421, CCR9-PE, Zombie-Aqua; HSPCs (excluding AX001): CD3-FITC, CD90-PE, CD49f-PECy5, CD38-PECy7, CD33-APC, CD19-A700, CD34-APC-Cy7, CD45RA-BV421, Zombie-Aqua; HSPCs (AX001): CD38-FITC, CD135-PE, CD34-PE-Cy7, CD90-APC, CD10-APC-Cy7, CD45RA-V450, Zombie-Aqua. Cells were either single-cell sorted for liquid culture into 96-well plates containing 50ul cell type-specific expansion medium, or (for AX001 HSPCs) bulk-sorted for MethoCult plate-base expansion.

### *In vitro* liquid culture expansion

We designed novel protocols to expand B and T lymphocytes from single cells into colonies of at least 30 cells. The B cell expansion medium was composed of 5ug/ml Anti-IgM (Stratech Scientific Ltd), 100ng/ml IL-2, 20ng/ml IL-4, and 50ng/ml IL-21 (PeproTech EC Ltd), 2.5ng/ml CD40L-HA (Bio-Techne Ltd) and 1.25ug/ml HA Tag (Bio-Techne Ltd), in Advanced RPMI 1640 Medium (ThermoFisher Scientific) with 10% fetal bovine serum (ThermoFisher Scientific), 1% penicillin/streptomycin (Sigma-Aldrich), and 1% L-glutamine (Sigma-Aldrich). The T cell expansion medium was composed of 12.5ul/ml ImmunoCult CD3/CD28 (STEMCELL Technologies) and 100ng/ml IL-2 and 5ng/ml IL-15 (PeproTech EC Ltd), in ImmunoCult-XF T Cell Expansion Medium (STEMCELL Technologies) with 5% fetal bovine serum (ThermoFisher Scientific) and 0.5% penicillin/streptomycin (Sigma-Aldrich). 25ul of fresh expansion medium was added to each culture every 3-4 days. Colonies (30-2000 cells per colony) were harvested either manually or robotically using a CellCelector (Automated Lab Solutions) approximately 14 days after sorting.

Sorted HSPCs from donors KX001, KX002, KX003 and CB001 were expanded from single cells into colonies of 200-100,000+ cells in Nunc 96 well flat-bottomed TC plates (ThermoFisher Scientific) containing 100uL of supplemented StemPro media (Stem Cell Technologies) (MEM media). MEM media contained StemPro Nutrients (0.035%) (Stem Cell Technologies), L-Glutamine (1%) (ThermoFisher Scientific), Penicillin-Streptomycin (1%) (ThermoFisher Scientific) and cytokines (SCF: 100ng/ml; FLT3: 20ng/ml; TPO: 100ng/ml; EPO: 3ng/ml; IL-6: 50ng/ml; IL-3: 10ng/ml; IL-11: 50ng/ml; GM-CSF: 20ng/ml; IL-2: 10ng/ml; IL-7: 20ng/ml; lipids: 50ng/ml) to promote differentiation towards Myeloid/Erythroid/Megakaryocyte (MEM) and NK lineages. Manual assessment of colony growth was made at 14 days. Colonies were topped up with an additional 50ul of MEM media on day 15 if the colony was ≥1/4 size of well. Following 21 +/- 2 days in culture, colonies were selected by size criteria. Colonies ≥ 3000 cells in size were harvested into a U bottomed 96 well plate (ThermoFisher Scientific). Plates were then centrifuged (500g/5min), media was discarded, and the cells were resuspended in 50ul PBS prior to freezing at −80C. Colonies less than 3000 cells but greater than 200 cells in size were harvested into 96 well skirted Lo Bind plates (Eppendorf) and centrifuged (800g/5min). Supernatant was removed to 5-10ul using an aspirator prior to DNA extraction on the fresh cell pellet. Sorted HSPCs from donor AX001 were plated onto CFC media MethoCult H4435 (STEMCELL Technologies) and colonies were picked following 24 days in culture.

### Whole genome sequencing of colonies

DNA was extracted from 717 colonies with Arcturus PicoPure DNA Extraction Kit (ThermoFisher Scientific), with the exception of larger HSPC colonies which were extracted using the DNeasy 96 blood and tissue plate kit (Qiagen) and then diluted to 1-5ng. DNA was used to make Illumina sequencing libraries using a custom low input protocol (*75*). We performed whole genome sequencing using 150bp paired-end sequencing reads on an Illumina XTen platform, to an average depth of 20x per colony. Sequence data were mapped to the human genome reference GRCh37d5 using the BWA-MEM algorithm (*76, 77*).

### Variant calling

We called all classes of variants using validated pipelines at the Wellcome Sanger Institute. Single nucleotide variants (SNVs) were called using the program CaVEMan (*78*), insertion/deletions (indels) using Pindel (*79*), structural variants (SVs) using BRASS (*80*) and copy number variants (CNVs) using ASCAT (*81*). In order to recover all mutations, including high frequency ones, we used an *in silico* sample produced from the reference genome rather than use a matched normal for the CaVEMan, Pindel, and BRASS analyses. Germline mutations were removed after variant calling (see below). For the ASCAT analysis we elected one colony (arbitrarily chosen) to serve as the matched normal.

Variants were filtered to remove false positives and germline variants. First, variants with a mean VAF greater than 40% across colonies of an individual were likely germline variants and were removed. To remove remaining germline variants and false positives, we exploited the fact that we have several, highly clonal samples per individual. We performed a beta-binomial test per variant per individual, retaining only SNVs and indels that were highly over-dispersed within an individual (*82*). For SNVs we also required that the variants be identified as significantly subclonal within an individual using the program Shearwater, and applied filters to remove artifacts resulting from the low-input library preparation. Detailed description of the artifact filters were provided previously (*75*) and the complete filtering pipeline is made available on GitHub (https://github.com/MathijsSanders/SangerLCMFiltering). For both the beta-binomial filter and the Shearwater filter we observed bimodal distributions separating the data into low and high confidence variants. We made use of this feature, using a valley-finding algorithm (R package *quantmod*) to determine the p-value cutoffs, per individual. We genotyped each colony for the set of filtered somatic SNVs and indels (per respective individual), calling a variant present if it had a minimum VAF of 20% and a minimum of two alternate reads in that colony.

We removed artifacts from the SV calls using AnnotateBRASS (https://github.com/MathijsSanders/AnnotateBRASS) with default settings. The full list of statistics calculated and post-hoc filtering strategy was described in detail previously (*83*). Somatic SVs were identified as those shared by less than 25% of the colonies within an individual. SVs and CNVs were both subsequently manually curated by visual inspection.

### Mutation burden analysis

We found that sequencing depth was a strong predictor of mutation burden in our samples. Therefore, in order to more accurately estimate the mutation burden for each colony, we corrected the number of SNVs or indels (corrected separately) by fitting an asymptotic regression (function *NLSstAsymptotic,* R package *stats*) to mutation burden as a function of sequencing depth per colony. For this correction we used HSPC genomes (excepting the tonsil samples, for which naive B and T cells were used), as lymphocyte genomes are more variable in mutation burden, and included additional unpublished HSPC genomes to increase the reliability of the model (Mitchell *et al.* in prep). Genomes with a mean sequencing depth of greater than 50x were omitted. The model parameters b0, b1, and lrc for each dataset for the model y = b0 + b1*(1-exp(-exp(lrc) * x)) are in Table S4. Mutation burden per colony was adjusted to a sequencing depth of 30.

We used a linear mixed effects model (function *lme*, R package *nlme*) to test for a significant linear relationship between mutation burden and age, and for an effect of cell subset on this relationship (separately for SNVs and indels). Number of mutations per colony was regressed on age of donor and cell type as fixed effects, with interaction between age and cell type, donor by cell type as a random effect, weighted by cell type, and with loglikelihood maximization.

### Mutational signature analysis

We characterized per-colony mutational profiles by estimating the proportion of known and novel mutational signatures present in each colony. For comparison, we included in the analysis 223 genomes from 7 blood cancer types: Burkitt lymphoma, follicular lymphoma, diffuse large B cell lymphoma, chronic lymphocytic leukemia (mutated), chronic lymphocytic leukemia (unmutated), and acute myeloid leukemia (*73*) and multiple myeloma (*51*). We identified mutational signatures present in the data by performing signature extraction with two programs, *SigProfiler (36*) and *hdp* (https://github.com/nicolaroberts/hdp). We used the *SigProfiler* denovo results for the suggested number of extracted signatures (12). *hdp* was run without any signatures as prior, with no specified grouping of the data. These programs identified the presence of 9 mutational signatures with strong similarity (cosine similarity >= 0.85) to Cosmic signatures SBS1, SBS5, SBS7a, SBS8, SBS9, SBS13, SBS17b, SBS18, SBS19 (version 3, (*36*)).

Both *SigProfiler* and *hdp* also identified the same novel signature (cosine similarity = 0.93), which we term the ‘blood signature’ or ‘SBSblood’. This signature is very similar to the mutational profile seen previously in HSPCs (*15, 16*). As the signature SBSblood co-occurs with SBS1 in HSPCs, leading to the potential for these signatures being merged into one signature, we further purified SBSblood by using the program *sigfit (84*) to call two signatures across our HSPC genomes - SBS1 and a novel signature - with the novel signature being the final SBSblood (Supplemental Figure 3, Table S5). SBSblood was highly similar to both the *hdp* and *SigProfiler* denovo extracted signatures (cosine similarity of 0.95 and 0.94, respectively) and had similarity to the Cosmic v3 SBS5 signature (cosine similarity = 0.87). One hypothesis is that SBSblood is the manifestation of SBS5 mutational processes in the blood cell environment.

We estimated the proportion of each of the 10 identified mutational signatures using the program *sigfit.* From these results we identified three signatures (SBS5, SBS13, SBS19) that were at nominal frequencies in the HSPC and lymphocyte genomes (less than 10% in each genome)-these were excluded from the analysis and the signature proportions were re-estimated in *sigfit* using the remaining 7 signatures: SBSblood, SBS1, SBS7a, SBS8, SBS9, SBS17b, SBS18 (Table S5).

### Ig receptor sequence analysis

In order to identify the immunoglobulin (Ig) rearrangements, productive CDR3 sequences, class-switch recombination and percent somatic hypermutation for each memory B cell, we ran *IgCaller* (*85*), using a genome from the same donor (HSPC or T cell) as a matched normal for germline variant removal. We considered the somatic hypermutation rate to be the number of variants in the productive IGHV gene divided by the gene length.

We estimated the number of mutations resulting from on-target (IGV gene) somatic hypermutation compared with those associated with SBS9. We first counted all IGV variants identified by Caveman pre-filtering, as we found that standard filtering removes many somatic hypermutation variants. We then estimated SBS9 burden as the proportion of SBS9 mutations per genome multiplied by the SNV burden. The SBS9 mutation rate per genome was the SBS9 burden divided by the ‘callable genome’ (genome size of 3.1Gb minus an average of 383Kb excluded from variant calling).

### Distribution of germinal center-associated mutations in B cells

We assessed the genomic distribution of the germinal center-associated mutational signatures, SBS9 and the SHM signature, in memory B cells. We performed per-Mb denovo signature analyses with *hdp* (no *a priori* signatures), treating mutations across all normal memory B cells within a given Mb window as a sample. The extracted ‘SHM’ signature (Table S5) had a cosine similarity of 0.96 to the spectrum of memory B cell mutations in the immunoglobulin gene regions, supporting the assumption that it is indeed the signature of SHM. In this analysis, SBSblood and SBS1 resolved as a single combined signature that we refer to in the genomic feature regression (below) as SBSblood/1.

We estimated the per-gene enrichment of SBS9 and SHM signatures across normal memory B and malignant B cell genomes (Burkitt lymphoma, follicular lymphoma, diffuse large B-cell lymphoma, chronic lymphocytic leukemia, and multiple myeloma). We first used *sigfit* to perform signature attribution of the signatures found in memory B cells (from the main signature analysis; SBSblood, SBS1, SBS8, SBS9, SBS17b, SBS18) and the extracted SHM signature from the above 1Mb *hdp* analysis, considering each 1Mb bin a sample. We subsequently calculated a signature attribution per variant. Gene coordinates were downloaded from UCSC (gencode.v30lift37.basic.annotation.geneonly.genename.bed). We calculated the mean attribution of variants in a given gene, representing the proportion of variants attributable to a given signature. We estimated the enrichment of SBS9 or SHM over genomic background per gene per cell type as the *p*-value of individual t-tests. While for this down-sampled dataset few genes were significant after multiple testing correction, analysis of full datasets with larger sample sizes show statistically significant enrichment in most presented genes (Figure 5) post-multiple testing correction (data not shown).

### Regression of SBS9 and genomic features

The *hdp* per-Mb memory B cell mutational signature results above were used to identify genomic features associated with the location of mutations attributable to a particular mutational signature. To achieve a finer-scale genomic resolution, each Mb bin was further divided up into 10Kb bins, and the proportion of each mutational signature in a Mb bin was used to calculate a signature attribution per 10Kb bin, based on the type and trinucleotide context of mutations in the 10Kb bin.

The number of mutations attributable to a particular mutational signature, per 10Kb window, was regressed on each of 36 genomic features (Table S3) (*64*). Noise was further removed from the replication timing data, using the GM12878 blood cell line data, and filtering the Wave Signal data by removing low Sum Signal (<95) regions, per Hansen *et al.* 2010 (*86*). SBS9 was analyzed separately from the SBSblood/1 combined signature. The number of mutations per signature per bin was calculated as the sum of the per-nucleotide probabilities per signature within a given bin. For the analysis of a given signature, a bin was only included if the average contribution of that signature was greater than 50%. This step ameliorates the problem of artificially high numbers of mutations being ascribed to a bin due to the combination of a trivially small attribution but a high overall mutation rate. This can occur in high SHM or SBS9 regions. This left 26,151 bins for SBS9 and 25,202 bins for SBSblood, out of 91,343 bins with mutations and 279,094 bins genome-wide. We also included a random sample of zero-mutation bins to equal 10% of the total bins.

We performed lasso-penalized general additive model regressions of the number of mutations per bin with the value of the genomic features. We used the *gamsel* function in R (package *gamsel*), with the lambda estimated from a 5-fold cross-validation of training data (2/3 the data). To estimate individual effect sizes, we performed general additive model regressions per genomic feature using the function *gam* (R package *mgcv*). The same analysis was also performed on HSPC mutations. The results for the full and individual regression models for each of SBS9 and SBSblood/1 in memory B cells and for all HSPC mutations can be found in Table S3.

### RAG and CSR motif analysis

We assessed the enrichment of V(D)J recombination (mediated by RAG) and class switch recombination (CSR, mediated by AID) associated motifs in regions proximal to lymphocyte SVs. We identified the presence of full length and heptamer RSS motifs associated with RAG binding and endonuclease activity (‘RAG motifs’) (*87*) for the 50bp flanking each SV breakpoint using the program FIMO (p<10^−4^) (*88*). Clusters of AGCT and TGCA repeats, associated with AID cytosine deamination and CSR (‘CSR motifs’), were identified in the 1000bp flanking each SV breakpoint using the program MCAST (p<0.1, max gap=100, *E*<10,000) (*89*). In order to estimate a genomic background rate of these motifs, we generated 100 genomic controls sets, randomly selected from regions of the genome not excluded from variant calling, and performed both the RAG and CSR motif analyses on these sets. The genomic background rate presented is the median of the 100 control datasets for each motif analysis. Both the RAG and CSR motif analyses were also performed for SVs from the PCAWG cancer genomes included in the mutational signatures analysis and for acute lymphoblastic leukemia genomes (*4*).

### Telomere length

We estimated the telomere length for HSPC and lymphocyte genomes (Table S2) using the program Telomerecat (*90*). Telomere lengths for all genomes for a given donor were estimated as a group.

### Timing of mutational processes

Following a procedure described previously (*59, 60*), we modelled the distribution of somatic mutations along the genome from the density of ChIP-sequencing reads using Random Forest regression in a 10-fold cross-validation setting and the LogCosh distance between observed and predicted profiles. Each mutation was attributed to the signature that most likely generated it and aggregated into 2,128 windows of 1Mb spanning ~2.1Gb of DNA. Signatures with an average number of mutations per window <1 were not evaluated due to lack of power. We determined the difference between models using a paired two-sided Wilcoxon test on the values from the ten-fold cross-validation. Epigenetic data were gathered from different sources (Table S6) (*61–63*) and consisted of 149 epigenomes representing 48 distinct blood cell types and differentiation stages and their replicates. Histone marks used included H3K27me3, H3K36me3, H3K4me1 and H3K9me3. To evaluate the specificity of SBS9 mutational profiles in memory B cells, we took the same number of mutations as in SBSblood with the highest association with SBS9 and compared models with an unpaired two-sided Wilcoxon test.

## Supplementary Figures

**Fig. S1.**
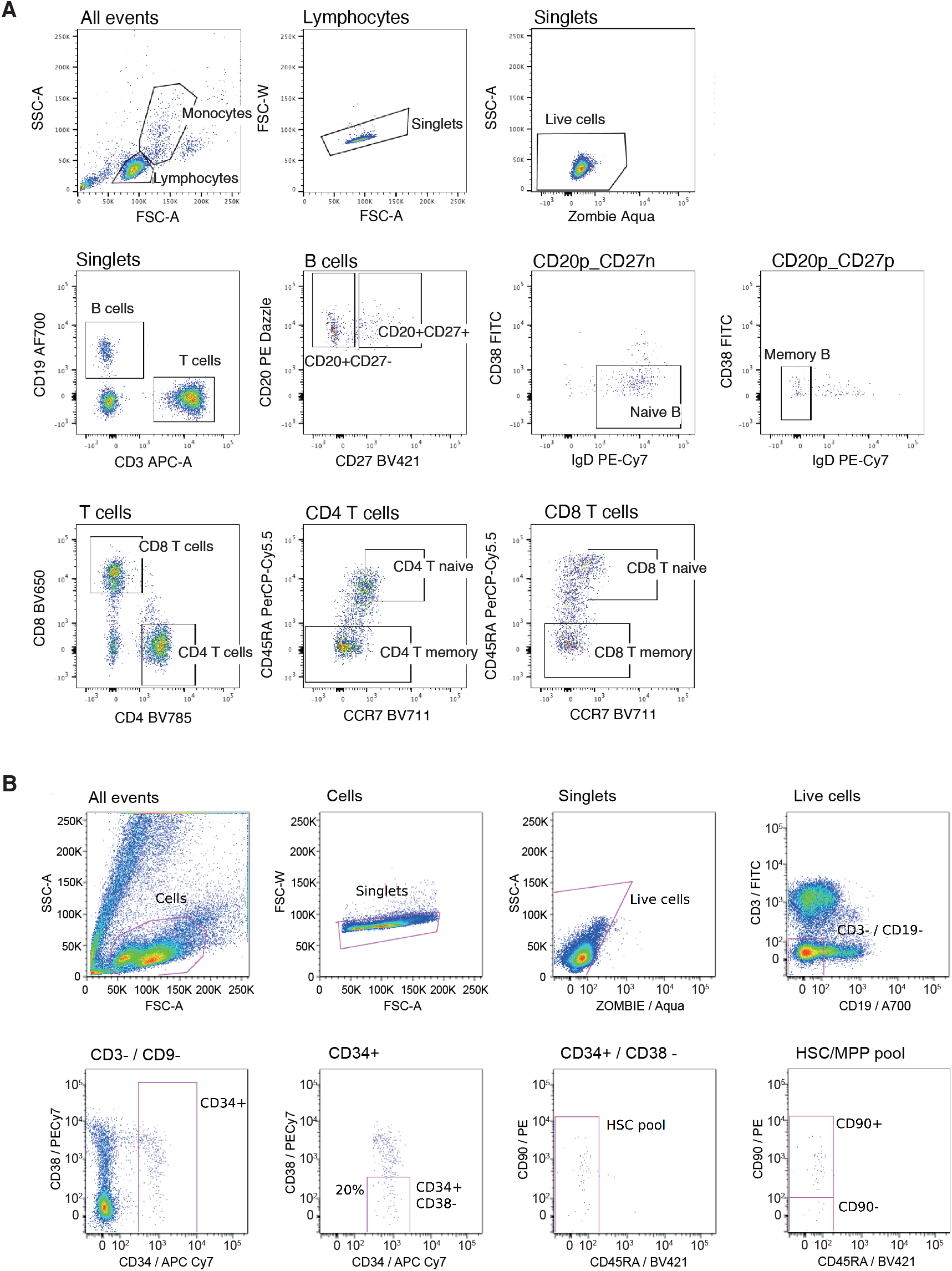
Flow cytometry gating for A) lymphocytes and B) HSPCs.

**Fig. S2.**
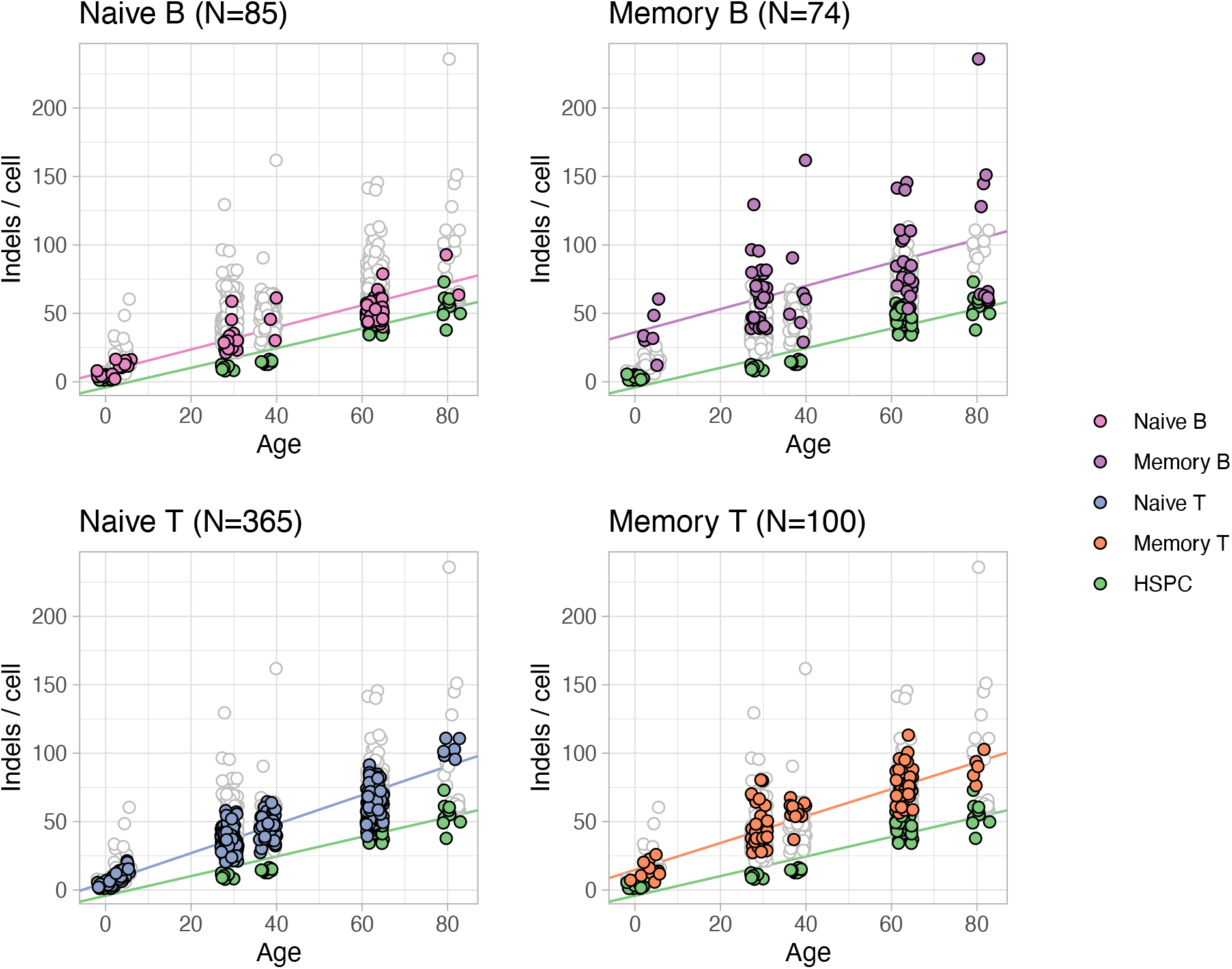
Indel mutation burden per genome for the four main lymphocyte subsets (colored points), compared with HSPCs (green points). Each panel has all genomes plotted underneath in white with grey outline.

**Fig. S3.**
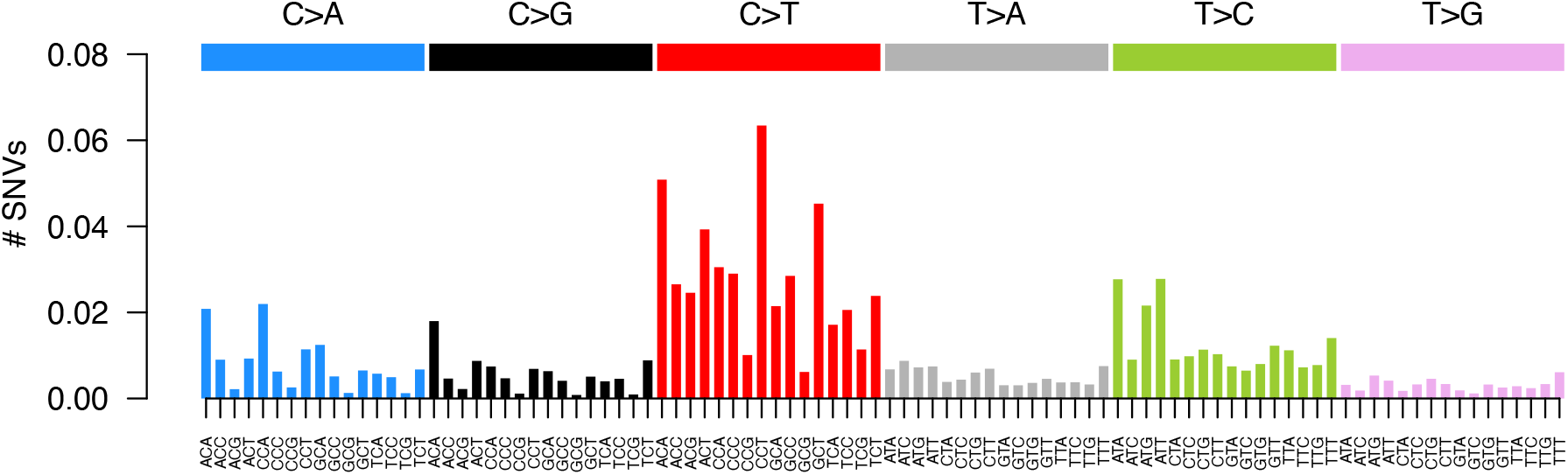
SBSblood signature identified using HSPC genomes and the program *sigfit* (excludes SBS1).

**Figure S4.**
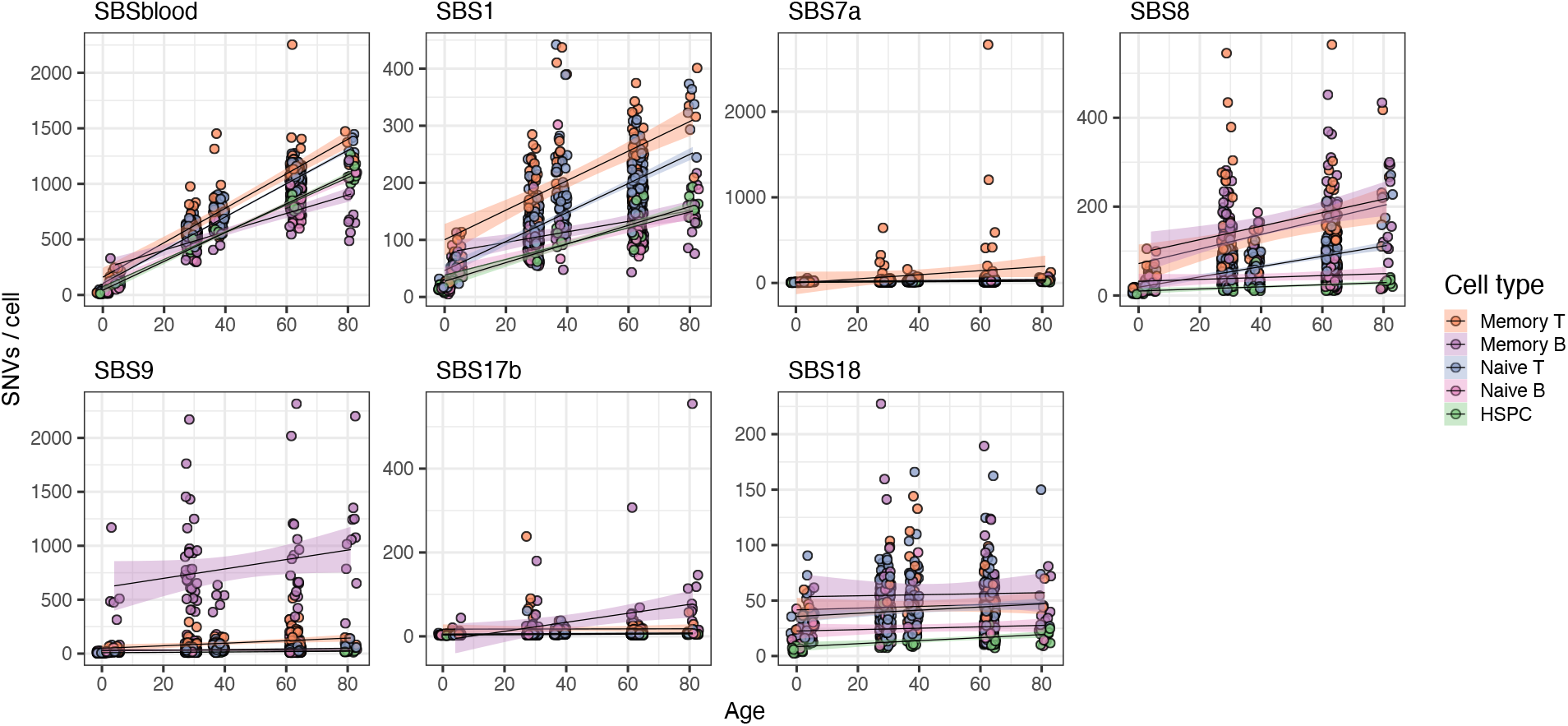
Mutation burden with age, per signature.

**Fig. S5.**
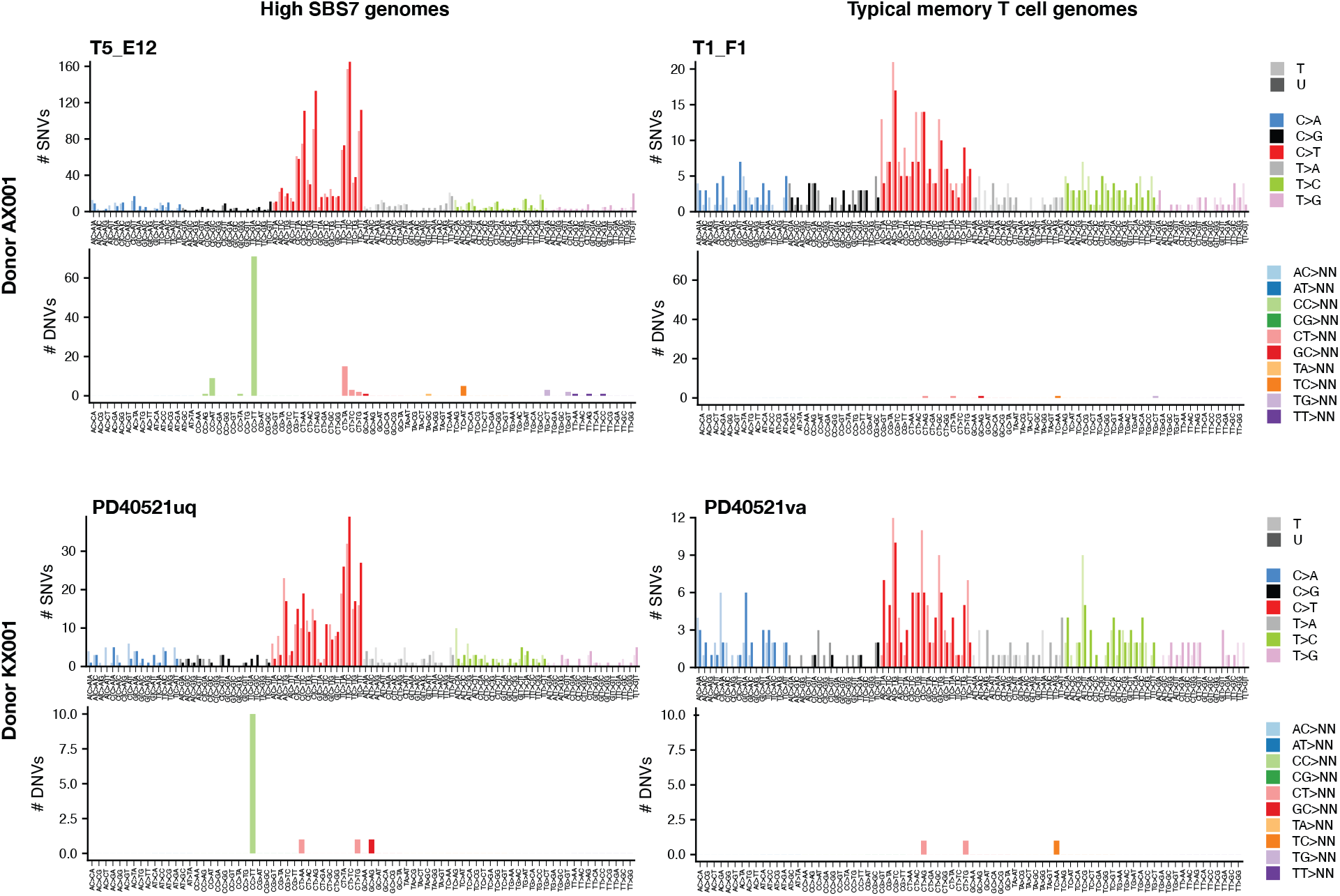
Single and dinucleotide mutational profiles for high UV cells. The SNV profile shows the transcriptional strand bias, with mutations on transcribed strands in a lighter shaded bar to the left and mutations on untranscribed strands in a darker shaded bar to the right. The two sets of plots on the left are of genomes with high levels of SBS7 (samples T5_E12 and PD40521uq), while the plots of the right are of genomes with the more common relative absence of SBS7.

**Fig. S6.**
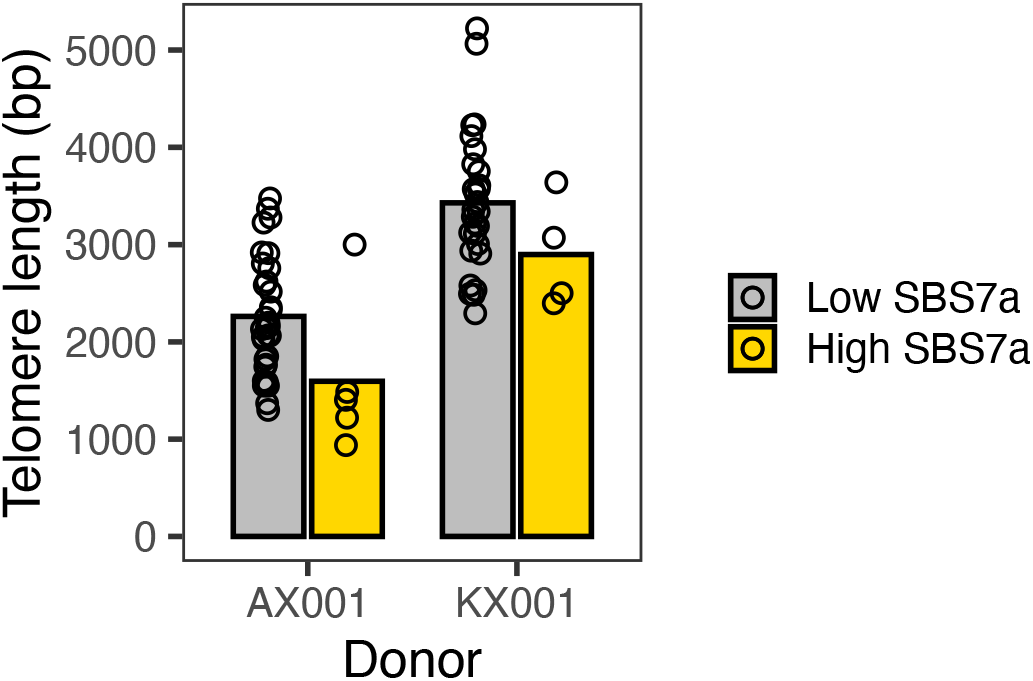
Telomere length in high UV signature memory T cells. A high UV signature memory T cell is defined as having greater than 9.5% (2 standard deviations above the mean) of its mutations attributable to SBS7a.

**Fig. S7.**
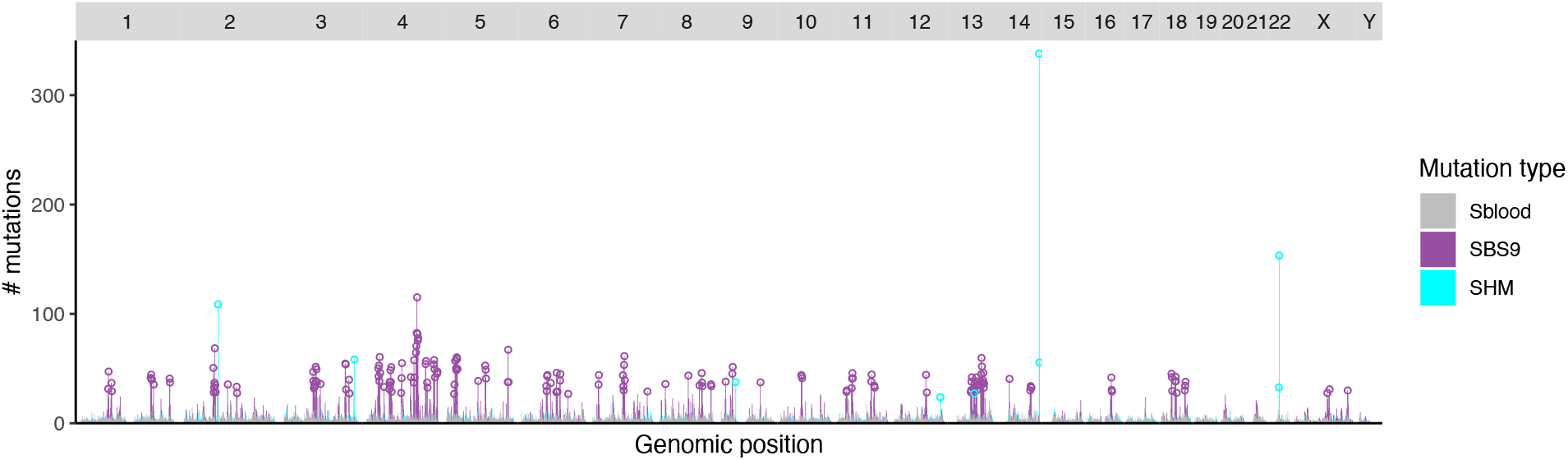
SBS9 and SHM signature genomic distribution in memory B cells. Signature extraction is performed per 1Mb genomic bin. Open circles denote bins with more mutations than 2 standard deviations above the mean.

**Fig. S8.**
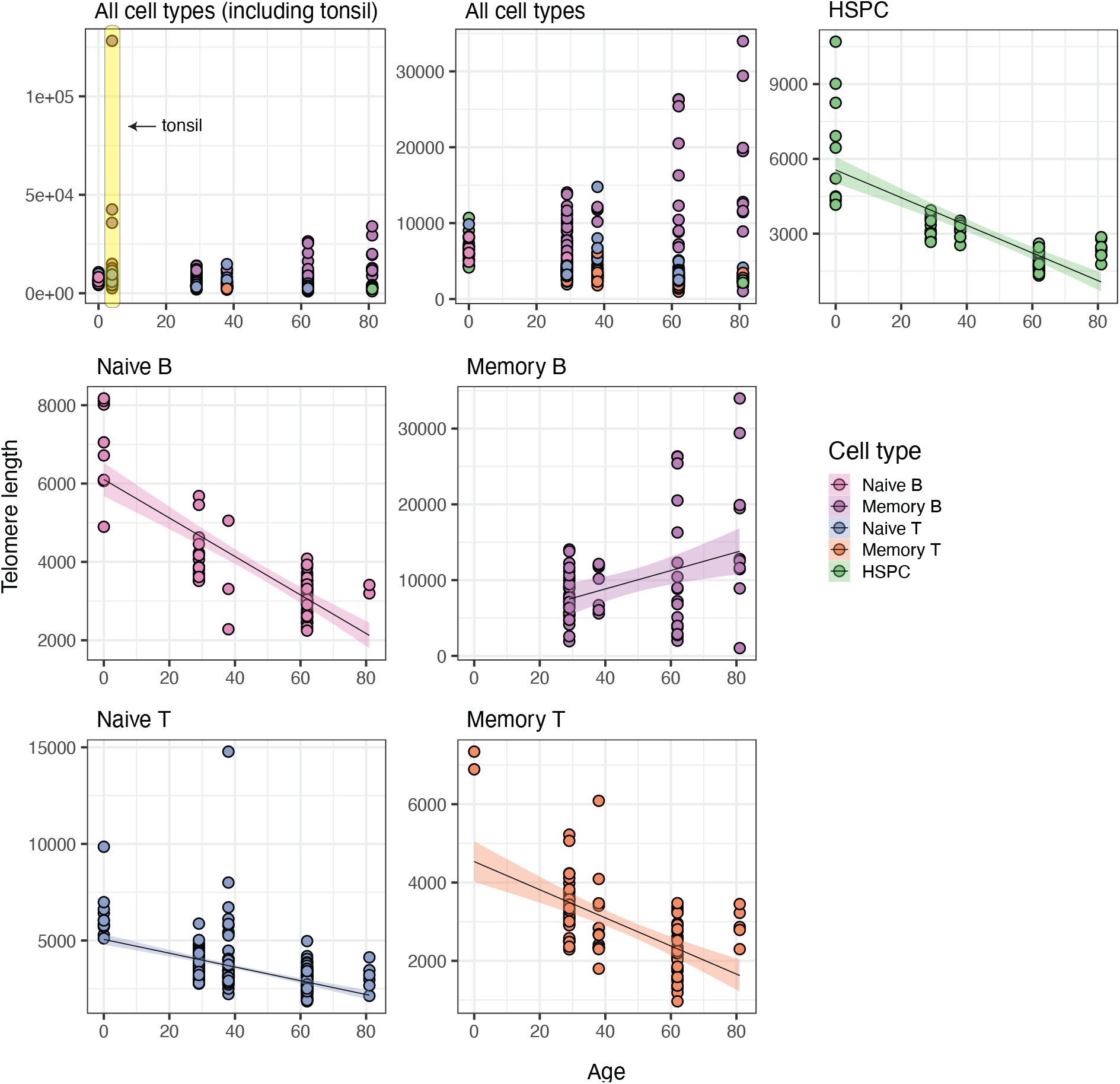
Telomere length as a function of age. The top left panel includes the tonsil-derived genomes, which have an exceptionally high variance in telomere length. The remaining panels exclude these genomes.

**Fig. S9.**
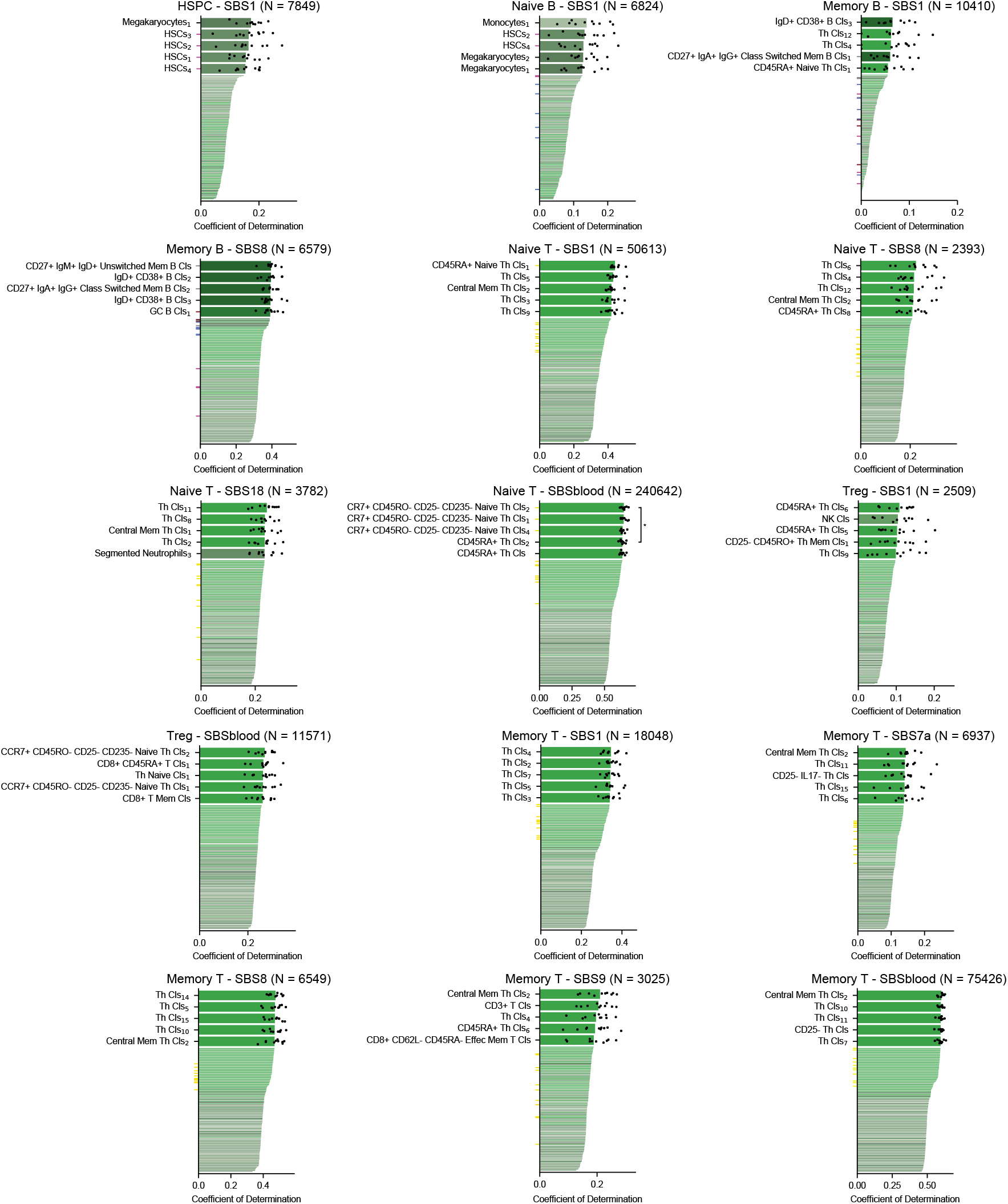
Performance of prediction of genome-wide mutational profiles attributable to particular mutational signatures from histone marks of 149 epigenomes representing distinct blood cell types and different phases of development (subscripts indicate replicates); ticks are colored according to the epigenetic cell type (purple, HSC; blue, naive B cell; grey, memory B cell; maroon, GC B cell); black points depict values from ten-fold cross validation; p-values were obtained for the comparison of the 10-fold cross validation values using the two-sided Wilcoxon test (Cls, cells; CS, class switched; GC, germinal center; HSC, hematopoietic stem cell; Mem, memory).

**Fig. S10.**
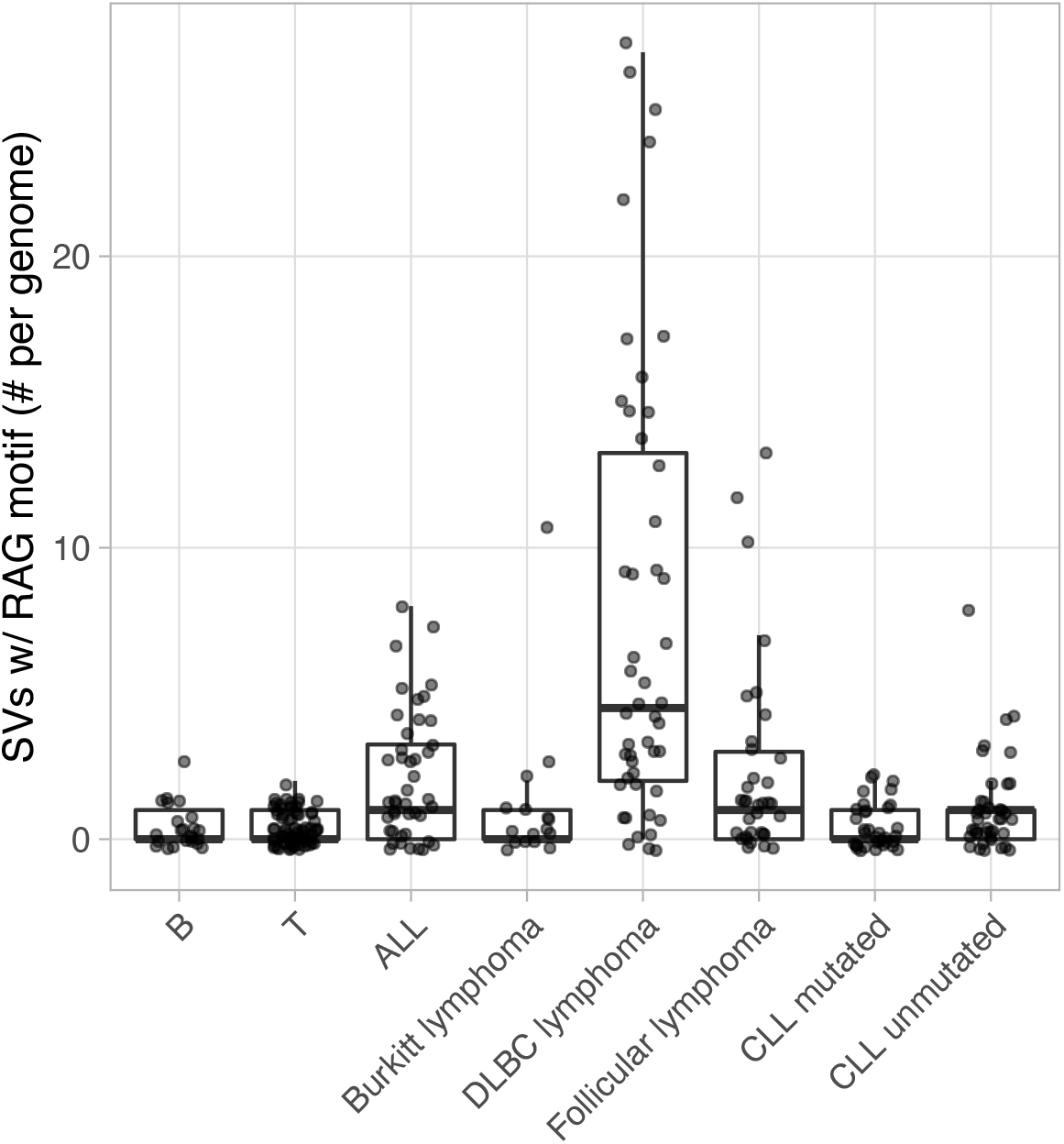
Number of SV’s with RSS (RAG) motifs within 50bp of a breakpoint.

